# The scotopic band: primate detailed scotopic vision and perceptual uncertainty

**DOI:** 10.1101/2022.01.25.477659

**Authors:** Shabtai Barash, Oleg Spivak, Peter Thier

## Abstract

Primates perceive detailed images in photopic (daylight) vision via a preferred-processing, high-acuity pathway, made up of a small photopic center (the fovea), and dedicated eye movements, whose function is to shift images of target objects to the fovea and keep them while foveal cones sense the image. Thus, the preferred processing pathway processes images serially, not in parallel. No such pathway is known for scotopic (night-light) vision or mesopic (twilight) vision – though details are informative in dim light, even vital; it remains unclear whether and how scotopic vision of details is accomplished. Here we show that primates do have a scotopic preferred-processing pathway. It consists of a ‘scotopic center’, located on a ‘scotopic band’, and eye movements that shift target images not to the fovea but directly to the scotopic center. In contrast to the stationary fovea, the scotopic center can relocate over the scotopic band. The scotopic center relocates to match changing visual conditions and contextual needs. Ambient light intensity is mapped monotonically onto the scotopic band, with mesopic vision at one end. The dimmer the light is, the more dorsal the scotopic center relocates on the scotopic band. Importantly, the scotopic center relocates (or ‘set’) not only in passive response to ambient light but also actively, driven by internal factors. In near-threshold conditions the scotopic center relocates to a more dorsal band location than in salient conditions. That the same relocation of the scotopic center can be instigated both by dimming the ambient light and by reducing the detectability of targets indicates that, at the core, the longitudinal axis of the scotopic band encodes the level of perceptual uncertainty.

## INTRODUCTION

High-acuity visual perception of primates is based on serial information processing. Rather than process the entire scene, one small image is processed after another. Acuity is constrained by the map of light-receptor density. Over the surface of the retina, the receptor density varies widely; only at the fovea, a small spot at central retina, is cone density high^1^. Hence, only the small image region that falls on the fovea can be viewed with high acuity. The term ‘direction of gaze’ refers to foveal gaze; it is determined by the position of the eyeballs in their sockets. As the eyes move, the instantaneous gaze direction moves with the eyes. Because they determine gaze direction, eye movements are crucial for high-acuity vision. The sensory impressions for high acuity vision accumulate through a series of fixations of ‘targets’, with quick jumps (‘saccades’) leading gaze from one target to the next. Vision evolves through fixations and saccades; consequently, fixation and saccadic eye movements turn photopic (daytime) high-acuity vision into a serial processing operation. What about scotopic (nighttime) vision – does it also incorporate a serial processing scheme based on some ‘scotopic center’, a confined retinal region of preferred scotopic processing? If it does, what is the retinal geometry of this location? Does it relate to the fovea? In what ways is it like the fovea? In what ways do the fovea and the presumed scotopic center differ, and why?

Three previous lines of research appear to support the possibility that a scotopic center does exist. The first line comprises observations about the geometry of rod density. Rods are dense dorsal to the fovea. Rod density systematically increases going dorsally, from the region just dorsal to the fovea, to a peak that was called either Dorsal Rod Peak (DRP) or Rod Hotspot^2,3^. Although the boundaries of the peak of rod density are not nearly as sharp as those of the foveal peak of cone density, the rod peak/hotspot is nevertheless a quite well-defined region, distinct from the rest of the retina. Rod density at the rod peak/hotspot is high; in fact, it is similar in order of magnitude to the foveal maximal cone density ^2,3^. Thus, at least the primary necessary condition for localized preferred scotopic processing appears to be satisfied.

The second line of research observed that, in darkness, monkeys direct their line of gaze above the target^4–6^. Could this ‘upshift’ reflect use of the rod peak? Namely, does the dorsal rod peak/rod hotspot functionally replace the fovea in darkness, in particular in scotopic vision? Although the idea initially seemed promising ^6^, problems confound this proposal. First, the size of the upshift does not match well the anatomical retinal distance between the fovea and the location of the rod peak. Moreover, the upshift varies largely from fixation to fixation, whereas the rod peak is fixed in its retinal place. Second, the upshift is observed in visual conditions that comprise dark background but are not scotopic – casting doubts on whether the upshift is indeed related to a retinal region dominated by rods. More specifically, the well-established time-course of dark adaptation^7,8^ reflects that rods remain saturated long (> 10 min) after dark onset. Upshift, on the other hand, was documented within seconds from dark onset^6,9^. In addition, published upshift studies generally lacked proper dark adaptation. Only with proper dark adaptation can the putative relationship of the upshift to scotopic vision be tested; preliminary evidence suggests that there might be upshift also in scotopic vision ^10^ but whether and how the upshift relates to scotopic vision, as compared to dark background in photopic vision, remains unclear. We will shortly suggest what this relationship might be.

The third observation which might support the existence of a scotopic center is that the pattern of saccades and fixations lingers on at night ^5,11^. Common thinking associates these movements solely with the fovea; presumably, a fixation keeps a target on the fovea, and a saccade subsequently shifts the next target to the fovea. The sensorimotor transformation of a saccade is thus presumed to reflect target foveation. The transformation activates the extraocular muscles so that the target’s image would fall on the fovea. This rationale is paradoxical for scotopic vision: cone-dense, the fovea is rod-sparse, hence insensitive in the dark. Why shift the image to an insensitive location? A putative scotopic center would constitute a solution to this paradox.

### Implanted search coils are accurate in scotopic conditions

In monkey studies of gaze direction, the technique of monitoring gaze direction using coils implanted over the sclera is now standard for more than 5 decades ^12,13^. Recordings with this technique are extremely stable and, critically, do not depend on the pupil’s contraction and dilation. Video-based eye tracking devices, now popular in studies of humans, is subject to artefacts and inaccuracies with the enlarged pupil of scotopic vision ^14–17^. Video-based tracking is especially perilous when studying a phenomenon such as upshift, which is itself a deviation of gaze direction. In addition, upshift is well characterized in monkeys (we have now recorded upshift in 14 monkeys), whereas reports in humans leave the presence of an upshift an open issue (see Discussion).

### Hypotheses

We propose the existence of a ‘scotopic band’: a confined, elongated retinal region extending dorsally from the immediate vicinity of the fovea. The anatomical work surveyed in the previous section suggests that, proceeding dorsally along the scotopic band (Fig 1A), cone density goes down, and rod density goes up (Fig 1A).

**Figure 1.**
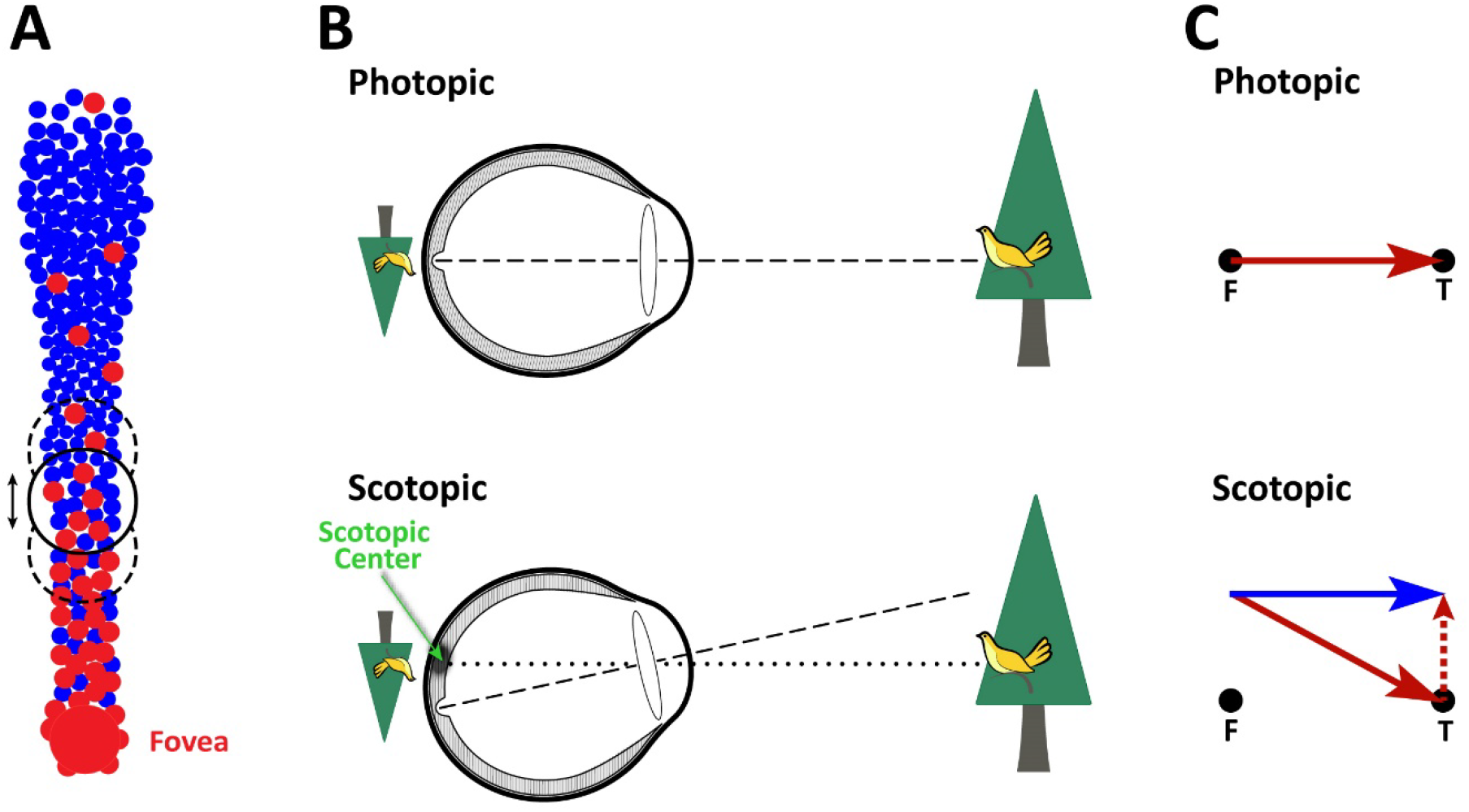
**(A)** Illustration of the hypothesis. Dorsal to the fovea there is a specialized ‘scotopic band’. At any time, a confined region within the scotopic band functions as a ‘scotopic center’. The target’s image is directed to the scotopic center, much as the target’s image is directed to the fovea in photopic vision. The scotopic center is the basis for scotopic serial processing, analogous to photopic high-acuity vision. The illustration schematically illustrates the photoreceptor density functions along the scotopic band, but this homology is only suggestive; our operational definition is purely functional. **(B)** Illustration of experimental prediction 1, scotopic upshift of fixation. In photopic vision, the target’s image (the bird on the tree in this schematic example) is fixated by the fovea. If in scotopic vision the target’s image falls on a scotopic center, an unsuspecting observer following only the foveal line of gaze would report that the subject looks above the target, not that the processing focus had switched from the fovea to the scotopic center. Such upshift was observed with dark background, but not scotopic. Dashed line, foveal line of gaze; dotted line, line of gaze to the presumed scotopic center. **(C)** Illustration of Prediction 2, saccadic sensorimotor transformations. The photopic sensorimotor transformation for saccades shifts the fovea directly from the initial fixation spot to the target. If all saccades foveate, including scotopic, then scotopic saccades too should end with the fovea on the target, even though they start with upshift (see panel B). Foveation (red arrows) is the ‘null prediction’. One alternate prediction is that the scotopic band setting, the location of the center on the band, would remain similar through the saccades. Thus, the saccades would start and end with upshift, reflecting a direct shift of the scotopic center from the fixation spot to the target (blue arrow).

We suggest, first, that at any instance, a current ‘scotopic center’ locus in the scotopic band carries a preferred role in scotopic vision much like the fovea does in photopic vision. Second, in scotopic vision, specialized sensorimotor transformations enable fixation and saccadic movements to shift the target’s image directly to the scotopic center and keep it there, much like the analogous sensorimotor transformations in photopic vision do with the fovea (Figs 1B,C; see ahead for details). Third, as visual and other conditions change, the active scotopic center moves along the scotopic band (black circles and arrows in Fig 1A). With these components, scotopic vision incorporates serial processing based on the scotopic center, much like the fovea-based serial processing of high-acuity photopic vision.

We further suggest that the scheme suggested here holds also for mesopic vision. Mesopic vision might be based more on the lower end of the scotopic band, in harmony with the fovea. However, in this manuscript we deal almost exclusively with the scotopic range.

### Operational definition

Our definitions of the scotopic center and scotopic band are pure functional, based solely on the eye-movement trajectories that are reported here. The accumulation of gaze directions while fixating a small fixation spot delineates the current center. We defer to the discussion questions such as how the scotopic band is related to the geometry of cone and rod density.

## RESULTS

### A scotopic center substitutes for the fovea

Is there indeed a scotopic center used to fixate targets in scotopic vision? We test a first experimental prediction related to fixations and a second experimental prediction related to saccades. We start with an example.

#### Directing gaze to the scotopic center

Imagine a monkey looking at a bird on a tree, first during daylight, then at night (Fig 1B). What would change at night, if the scotopic center hypothesis is valid? The eye would rotate, so that the bird’s image would fall on the scotopic center rather than on the fovea (dotted line, lower sketch of Fig 1B). Consequently, the fovea’s line of gaze would now be directed above the bird (dashed line, lower sketch of Fig 1B). Because measurements of gaze direction refer to the foveal line of gaze, the rotation appears to be an upshift of gaze direction, much like that observed in non-scotopic darkness ^4,6,9^. At first sight, it could seem strange that the bird’s image falls away from the fovea. However, this situation could simply reflect the switching from photopic to scotopic state. Thus, the bird’s image could simply fall on the scotopic center, which is located above the fovea. Thus, according to the scotopic center hypothesis, the visual system’s serial processing switches from using the fovea as its preferred retinal region to using the scotopic center.

Thus, the first experimental prediction transpiring from the hypothesis is that in scotopic vision there is upshift. Because the term upshift directly describes the result, we would use it, but the connotation is always that of the implicated switching from the fovea to the scotopic center.

We studied 3 rhesus monkeys already trained in visual and oculomotor tasks. A daily session started with a standard calibration procedure, conducted in bright light. Then followed: a photopic study, a 45-min dark-adaptation interval in complete darkness, and a scotopic study. In trials of both photopic and scotopic studies, the monkey had to fixate a central fixation spot, then follow with his eyes the spot’s jump to another location and fixate the target at its new location. The initial fixation spot and all targets were on the horizontal meridian. We collected data in photopic and scotopic conditions; we will call data – trials, saccades, and so on – ‘photopic’ or ‘scotopic’ according to the conditions that prevailed when collected. Photopic data will be displayed in red, scotopic in blue.

We compared the mean gaze directions in photopic and scotopic trials. We made 2 sets of comparisons. The first set compared fixation of the central target that preceded the saccades. The second comparison encompassed fixations of the peripheral targets, during the post-saccadic fixations. The comparisons are illustrated in Fig 2A, in the left and right panels for each monkey, respectively. Each dot in the scatterplots reflects the mean gaze direction of one trial. The dots are positioned in the scatterplots according to the gaze direction of the individual trial. To be clear, the measurements reflect the direction of the foveal line of gaze, regardless of whether the fovea actively senses (as in photopic vision) or not (as in scotopic). The mean gaze directions were assessed from 0.25-s intervals, in the post-saccadic data long enough after preceding correction saccades to have ended. Thus, both photopic and scotopic data reflect fixations of targets at the same locations, all on the horizontal midline of the screen. Nonetheless, foveal gaze directs upwards in scotopic vision; the clusters of blue dots are positioned above the red clusters in all 6 panels. No systematic horizontal shift is present. The Figure illustrates results of many trials (847, 594, 398 photopic trials, 1216, 611, 588 scotopic, correspondingly in M1, M2, and M3). Nonetheless, the blue and red clouds are almost perfectly separate. The only exception is a small set of scotopic trials (∼60) of monkey M1, in which the foveal line-of-gaze directs to the target even though M1 is using scotopic vision. We will allude to this small set later. The histograms of the vertical components of gaze direction, for photopic and for scotopic trials, are illustrated adjacent to the respective scatterplots. Note that the axes of the vertical component of gaze direction are common to the scatterplots and the histograms. The histograms of the vertical components reflect the almost perfect separation of photopic and scotopic gaze directions. The overlap of the histogram is very small, 0 in M1 (not counting the 60 trials mentioned above), <4% (23/594) in M2, 0 in M3. In all 3 monkeys, both at the center and in the periphery, the means of the photopic and scotopic foveal-line-of-gaze directions differ from each other (*p<10*^*-264*^ in each comparison, t-tests). In line with previous studies, we define upshift to be the difference of the vertical components of mean scotopic position from the mean photopic. Applying this definition, the upshift is 6.2 deg in M1, 2.1 deg in M2, and 5.6 deg in M3. Thus, the prediction stated at the start of this section is validated. There is upshift not only briefly after darkness onset, but also long after darkness onset, in scotopic vision.

**Figure 2.**
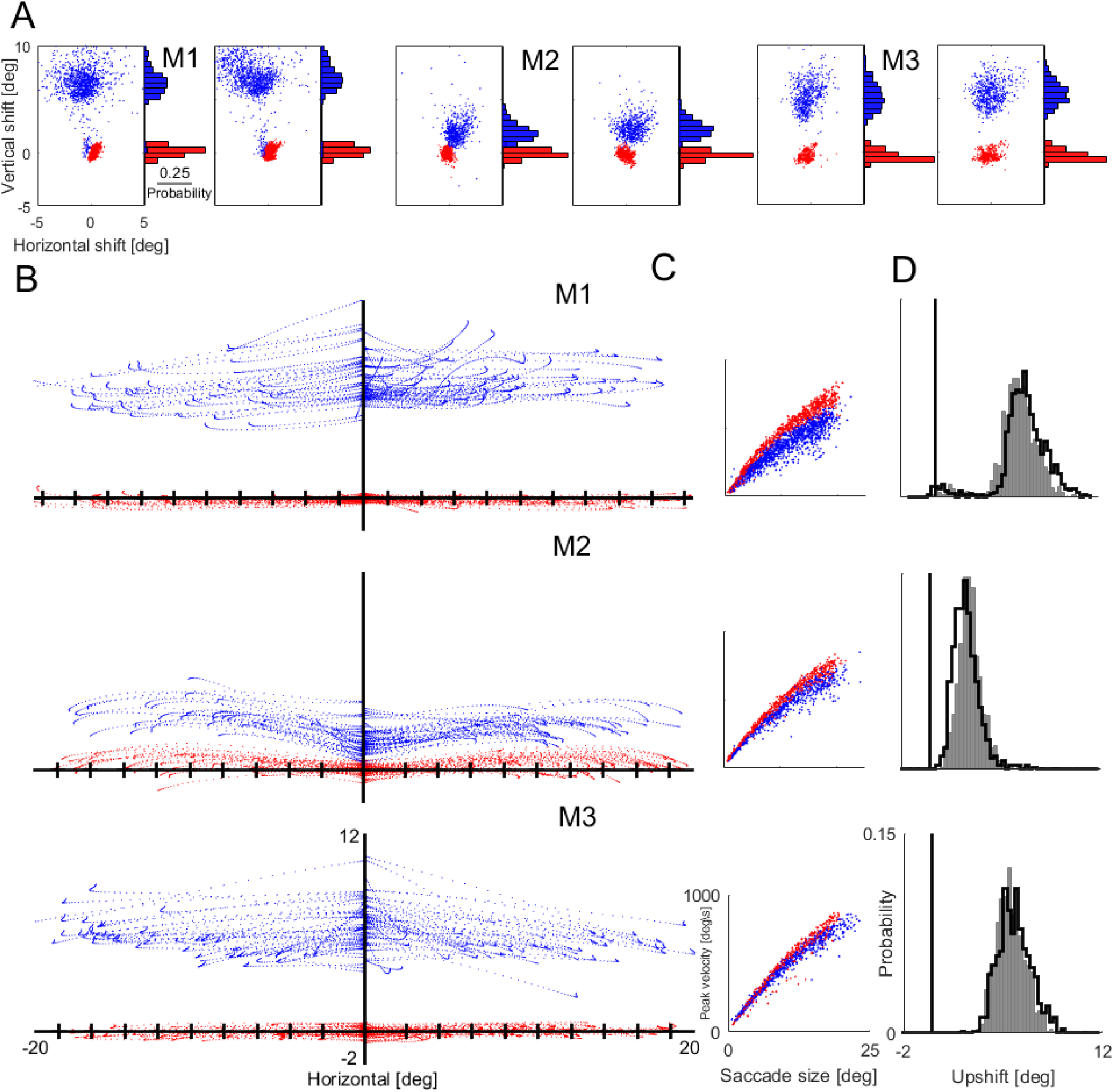

The blue clusters in Fig 4A are larger than the red clusters, in all 6 panels. The blue clusters effectively chart the extent of the scotopic center. The blue clusters are larger than the red clusters; this observation indicates that the scotopic center is larger than the fovea. These results are consistent with reported variation of photopic upshift ^6,9^.

The upshift for targets on the horizontal meridian illustrated in Fig 2A indicates that upshift is indeed present in scotopic conditions, not only in the photopic phase of dark adaptation in which upshift was previously documented. That the presence of upshift is not limited to the horizontal meridian is indicated by the studies of upshift in photopic dark ^6^ as well as by preliminary results in scotopic conditions ^10^.

#### Saccades targeting the Scotopic Center

Schematic Fig 1C illustrates the sensorimotor transformations of saccadic eye movements, in photopic vision (top) and scotopic (bottom). The sensorimotor transformation of a typical photopic saccade is marked by a trajectory that leads the fovea’s line of gaze directly from the initial fixation spot to the target (red arrow in Fig 1C, top panel). What happens in scotopic vision? The eye begins with an upshift, as seen in Fig 2A. Hence, for the line of foveal gaze, the saccade’s starting point is drawn in Fig 1C as the point at which the blue and red arrows initiate, right above the fixation spot (marked ‘F’). Now, if, in line with standard thinking, all saccades always aim to foveate, then we expect the saccade to end with the fovea directed to the target; in other words, with the foveal line of gaze on the target. The target is marked in the lower panel of Fig 1C by the black spot marked ‘T’, and the expected movement by the red arrow. Thus, by standard thinking, we expect no upshift to be present when the saccade comes to its end. Because we do expect eventually upshift to occur, we would expect it to build up after the saccade (dotted red line in Fig 1C). Thus, the foveation hypothesis, common in standard thinking, leads to the experimental prediction just described, depicted in Fig 1C’s lower panel in red. In the present study, this is the null prediction. Transpiring from the hypothesis that a scotopic center is used in scotopic vision, much like the fovea in photopic vision, is the alternate prediction – that saccades shift the target’s image *directly* to the scotopic center. By the scotopic center hypothesis, when the saccade begins, the fixation spot’s image falls on the scotopic center; and when the saccade ends, the target’s image falls on the scotopic center. Consequently, the foveal line of gaze would be directed above T at the end of the saccade, just as it was directed above F at the onset of the saccade. Because in the present study both F and T are on the horizontal meridian, we expect the saccade (blue arrow in Fig 1C) to be close to horizontal. Furthermore, we expect the saccade to end clearly above T.

Fig 2B illustrates trajectories of example saccades made by the 3 studied monkeys, photopic saccades (red traces) and scotopic saccades (blue traces). All trajectories were horizontally shifted to start from the vertical meridian (this shift was applied only to the example traces of Fig 2B, not to the analysis of the entire database in Figs 2C,D). The superimposed photopic (red) trajectories remain close to the horizontal axis; these saccades took a largely straight course, ending with the fovea directed close to the relevant trial’s target. The scotopic trajectories (blue) are very different: as expected, the foveal line of gaze starts directed above the central fixation spot. The vertical range of the scotopic gaze directions is larger than the vertical range of the photopic gaze directions, in line with the observation of Fig 2A, likely reflecting, again, that the scotopic center is indeed larger than the fovea. After the saccades start, the trajectories of the foveal line of gaze transpire in a near-horizontal direction. A few traces have a slight downward component; others go slightly up. Importantly, the saccade endpoints remain in the vertical range of the starting points, suggesting that the saccades end with the target’s image falling in the active scotopic center. Turning to the predictions sketched in Fig 1C, the example saccade trajectories are similar to the horizontal blue arrow of Fig 1C. This similarity supports the alternate prediction, that the saccades shift the scotopic center directly to the target. The trajectories are strikingly inconsistent with the null prediction. Not even one saccade illustrated by a blue trajectory ends with the fovea directed at the target. The endpoints of the blue trajectories are all above the endpoints of the red trajectories. Thus the example saccades suggest that the null prediction is to be rejected.

Figs 2C,D show the results of a systematic analysis of all saccades recorded in each monkey in this experiment. Fig 2C confirms that scotopic saccades show a main sequence similar to that of photopic saccades. The speeds of scotopic saccades in our data were somewhat slower than the speeds of photopic saccades, but we do not know if this would have generalized to all experimental conditions; in any case, the blue main sequence plots appear to be standard saccade main sequences. The main results are illustrated in Fig 2D. These data make it possible to test the null prediction and evaluate the alternate prediction systematically, based on the entire database. Fig 2D shows, for each monkey, two histograms of upshift values from single trials. The filled gray histograms show the upshift during fixation of the central fixation spot (‘F’ in Fig 1C), just before saccade onset; the unfilled histograms, depicted by the solid line, show the values of the upshift at the peripheral target locations, briefly after the saccade offsets. Zero upshift, that is, no upshift, is marked as a vertical line on each panel of Fig 2D. Let’s now assess the predictions.

The null prediction states that that in scotopic vision, as in photopic, saccades shift the foveas to the target. Thus, at the end of the saccade there should be no upshift. In other words, the trial-by-trial upshift histogram should be centered on 0 upshift. This prediction is decisively rejected. The means of the end-saccade upshift histograms are positive for all monkeys, with p=0 (t-tests). For M2 and M3 no trials start or end close to 0; for M1, only a tiny minority (4.8 %) do not show up upshift. These trials probably reflect the same rare state as the 60 fixation trials mentioned above. Thus, the null prediction is decisively rejected.

Rejecting the foveation hypothesis is the main result of this section. Nonetheless, for evaluating the alternate prediction, we compare the start-saccade and end-saccade histograms of each monkey. The start and end saccade histograms are not identical – the two-sample Kolmogorov Smirnov test gives *p<10*^*-6*^, for each monkey. The means differ, too – but only by little: on going from start-to end-saccade, the mean upshift values change by 8% (M1), -9% (M2), 5% (M3). This change is but a small fraction of the size of the scotopic center. Thus, the saccade appears to shift the target to the active scotopic center, much as photopic saccades shift the target’s image to the fovea. After all, photopic saccades do not end with the target always at the same region of the fovea.

We set out to study saccades over the horizontal meridian with the hope that horizontal target shifts would provide a clear-cut, visible test of the foveation hypothesis, and they did (Fig 2B,D). We may still ask if a larger set of target positions would have yielded a different answer. With regard to the main point, of testing the foveation hypothesis, a larger set wouldn’t have mattered: rejecting a prediction for the subset of horizontal saccades shows that the foveation hypothesis fails. Upshift far below or far above the horizontal meridian changes nonlinearly ^6,9^ as do saccades to these areas ^9,18^ so the question remains open if the relationship between static upshifts and saccades is less disciplined far from the horizontal meridian, though limited evidence suggests this does not happen ^9,19^. In summary, the foveation hypothesis is rejected; the scotopic center hypothesis is strongly supported close to the horizontal meridian and appears to hold also away from the meridian.

### The Scotopic Band

#### Introduction

By now, we have presented evidence for the existence of a scotopic center and of eye movements tailored to enable its use. Now we get to the second part of the hypothesis guiding this manuscript. We suggest that, as visual conditions change, the position of the scotopic center changes too. We define the scotopic band as the range of positions taken by the scotopic center. The results that we will present indicate that the scotopic band is an elongated structure positioned on the vertical meridian. The main axis of the scotopic band is near-vertical. The ventral end of the scotopic band is near the fovea, just dorsal to the fovea. We know less about the dorsal end of the scotopic band, and it might differ from monkey to monkey, but it appears that typically the scotopic band extends at least to the representation of an eccentricity of 10 - 15 deg visual angle.

Photopic vision does well with a small, near circular photopic center (the fovea). Why would scotopic vision require an elongated band, over which the scotopic center would relocate? Perhaps the reason has to do with the availability of information in the photon flux that falls on the retinas in natural conditions. Photopic vision remains largely unchanged over its operational range because photons are dense enough to maintain enough information about objects large enough to be of interest for vision throughout the photopic range. Scotopic vision at the bright end of the scotopic range might be analogous to photopic vision. However, over the scotopic range the situation changes. With dimmer ambient light, there is less and less information available because there are not enough photons. So, at the dim end of the scotopic range the required processing is very different from that of photopic vision. This might be related to the change in rod (and cone) density over the scotopic band ^2,3,20^.

Although the experiments of this paper address almost exclusively scotopic vision, the scotopic band might encompass also mesopic vision. The two might be integrated over the scotopic band.

#### Background luminance impacts the scotopic center’s position on the band

On studying the upshift, we were struck by incidental observations suggesting that the background luminance, reflecting in nature the intensity of the ambient light, affects the size of the upshift. This observation, together with the high variability of the upshift, led us to suggest the notion of the scotopic band. Setting out to systematically test whether a scotopic band might indeed be present, we started by assessing the upshift for a range of background luminance levels. We start by illustrating the effect of background luminance on the upshift of single trials, in an example block sequence (Fig 3A). We kept the target very small, with fixed luminance; because the target remains unchanged, observed changes in the upshift reflect the changed background, not the fixed target ^6,9^. The monkey was dark-adapted for 45 minutes. Then testing commenced. A sequence of brief blocks followed each other, with background luminance maintained during a block and increased slightly from block to block. Each dot reflects the upshift of a single trial. All trials of a block are drawn with the same color. The squares mark each block’s mean upshift values with each square’s color the same as that of the trials of the block.

**Figure 3.**
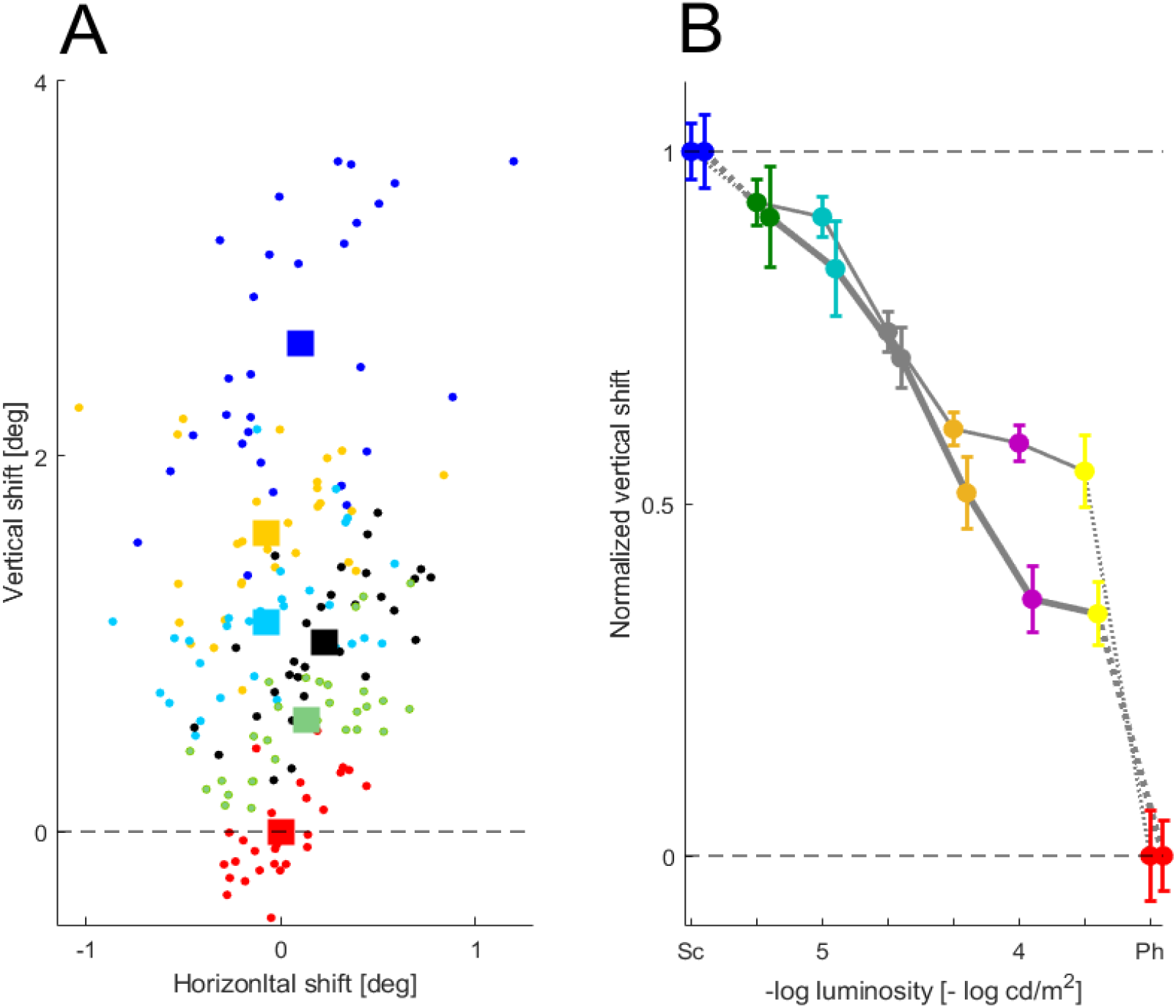
Evidence for the existence of the scotopic band: the darker the background, the greater the upshift is. **(A)** Example session, with trial-by-trial gaze direction data for background luminance increasing from block to block, from dark to photopic level. Every dot illustrates foveal gaze direction in one trial, while fixating small (0.02 deg radius) bright (60 cd/m^2^) targets. The plot depicts six consecutive blocks, of 30 trials each. Every block had its own background illumination: 10^−6^ cd/m^2^ (blue), 10^−5^ cd/m^2^ (yellow), 10^−4^ cd/m^2^ (cyan), 10^−3^ cd/m^2^ (black), 10^−2^ cd/m^2^ (green), 7 cd/m^2^ (red). The squares depict the mean foveal gaze direction in each block. The dashed line marks zero upshift (no upshift). **(B)** Mean upshift for several levels of background luminance. Data were collected from 2 daily sessions for each monkey, one block from each session, and normalized to each monkey’s scotopic upshift. Monkey M2’s data are shown with the thick grey line, M3’s as thin grey line. Each point shows the mean normalized shift of the 2 blocks with the standard error. Full scotopic shift is shown in blue, photopic (7 cd/m2 background) in red; the intermediate points show background levels in the scotopic range, green: 0.4*10^−5^, cyan: 10^−5^, grey: 0.2*10^−4^, orange: 0.4*10^−4^, purple: 10^−4^, yellow: 2*10^−4^ cd/m^2^). Target luminance was 10^−3^ cd/m^2^, target radius: 0.25 deg. The dashed vertical lines mark vertical zero (no upshift) and 1 (maximal upshift).

The scotopic range is covered in about 4 blocks (blue, yellow, cyan, black, with background luminance stepped up from block to block). Black reflects luminance at the transition between scotopic and mesopic (see Legend and Methods). Green reflects an even more intense background; red is full photopic.

The upshift varies considerably from trial to trial. Nonetheless, the mean upshift values show a robust trend: as the background luminance increases, the upshift decreases, gradually, from block to block. Although most of the decrease is in the deep scotopic range (blue to cyan), some of the decrease is present well into the mesopic range. This is consistent with the thinking that the scotopic band is involved in both mesopic and scotopic vision, and they form a continuum.

Guided by the example session illustrated in Fig 3A, we turn to the first experimental prediction emerging out of the scotopic band hypothesis: as the background luminance increases within the scotopic range, the prediction suggests that the upshift decreases. We tested 2 monkeys, M2 and M3. The results are illustrated in Fig 3B. The upshift of the two monkeys was normalized, to help compare their data. The normalization was linear and defined so that the photopic upshift (that is, no upshift) was set to 0 and the full scotopic upshift was set to 1. The entire scotopic range was sampled. Targets were small (0.25 deg radius) and bright enough to be scotopic, yet salient at all tested background luminance levels (10^−3^ cd/m^2^) (see next section). Thus, Fig 3B illustrates the upshift at several luminance values throughout the scotopic luminance range. Importantly, the monkeys were fully dark adapted to the level of the tested background throughout data collection.

In both monkeys, the upshift goes down with increasing background luminance. The normalized graphs of the upshift of the 2 monkeys are both monotonically decreasing and similar to each other (blue to violet dots, up to 10^−4^ cd/m^2^).

This result strongly supports the notion of a scotopic band. As the background stepped up, from block to block, throughout the scotopic range, the scotopic center systematically relocated to more ventral locations, closer to the fovea. The range of these movements of the scotopic center makes up the scotopic band.

### The scotopic band near threshold

#### Near-threshold fixations

The existence of the scotopic band implies that using the scotopic center necessitates repeated acts of choosing where on the band to position the active scotopic center. We will call this act of choosing a desired position on the band “scotopic band setting”. An effect of scotopic band setting is that the sensorimotor transformations involved in scotopic saccades and fixations are more complex than those of photopic vision, because the scotopic transformations depend on a parameter (position on the band). Moreover, the question comes up: does scotopic band setting happen always automatically? The previous case of scotopic band setting, by the background luminance (Fig 3), can, it appears, happen as automatically as, say, photoreceptor adaptation. Are there other cases of scotopic band setting that might not appear to be automatic? Guided by this question, we turn to study threshold situations.

Large targets allow fixation accuracy to be measured only roughly, because adequate fixation can aim at any location within the large target. To avoid this caveat we have used in the studies described so far only small targets; to ensure that these small targets are salient, the targets had to be bright, high on the scotopic luminance scale. We now seek to explore fixations of dimmer, larger targets; such targets are common in scotopic vision. Does their fixation involve upshift? What part of the scotopic band is used? Of particular interest are threshold situations. Before we turn to upshift, we will ask: what parameters determine thresholds for making saccades to scotopic targets? Then we will go back to upshift and ask: does a threshold situation influence the scotopic band setting? That is, does upshift of fixation near threshold differ from the upshift of a physically similar target while it is salient? Such finding would suggest that while close to threshold, scotopic band setting is influenced by a non-automatic process.

The rationale of studying threshold situations is the following: Light conveys information. As luminance goes down, photons become less ubiquitous, the information carried by the light changes. The ambient light significantly influences the image’s statistics. Near threshold, noise is especially high. The visual system must respond to this change of available information. Does the signal to noise ratio influence the monkey’s setting of the scotopic band?

The following study aimed at taking on these questions. Two monkeys took part in this study. Example sessions of the 2 monkeys are illustrated in Figs 4A,B, which will be described shortly; altogether we ran 5 sessions of M3, then 7 sessions of M2. The results will be illustrated by a single session for each monkey (Figs 4A,B); all the analysis that will be presented pertain to the entire database of all sessions recorded in this study.

**Figure 4.**
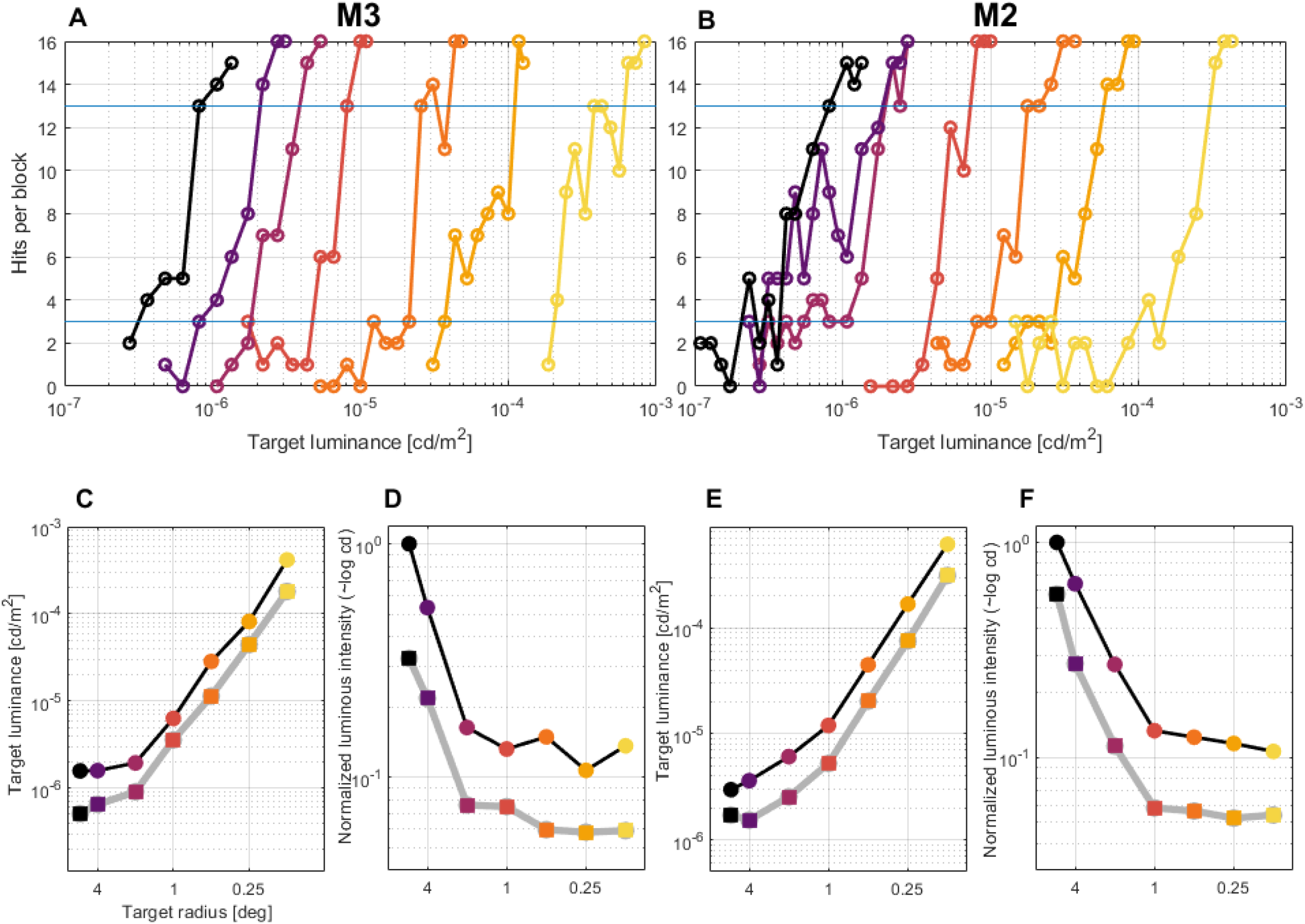
The monkeys’ performance in a study of visual fixations in near-threshold versus salient conditions. **(A)** Example of a single session of monkey M3. Each dot reflects the number of hits in a 16-trial block using a target of the specified luminance. The session is made up of block sequences, color coded; each sequence uses a fixed target size. While keeping target size fixed, in each consecutive block the target is made slightly brighter. The resulting psychometric functions reflect the rise from no response (0-2 hits, under the lower blue horizontal line) through threshold (3-13 hits, between the 2 horizontal blue lines) to salient, full performance (14-16 hits, above higher blue line). Target radii in visual degrees, from left: 5.5, 4, 2, 1, ½, ¼, 1/8. **(B)** Same as (A), for monkey M2. **(C)** Mean target luminance required for reaching threshold (gray line) and salience (black line) for each target size, computed from all sessions of M3. The smaller the target, the more luminous the target must be for reaching performance. **(D)** M3’s performance analyzed as in terms of normalized luminous intensity, proportional to target luminance times target area, also the total number of photos absorbed per unit time. Small targets need about the same level of luminous intensity to reach threshold and salience, almost invariant of target size. Large targets require graudally more luminous intensity. **(E)** and **(F)** Analysis of monkey M2’s performance; same format as **(C)** and **(D)**.

For creating a threshold situation that would allow assessing the effect on the upshift, we varied (a) the target’s size, and (b) the target’s luminance. Figs 4A,B illustrate the results of an example session of each of the participating monkeys. A session was made up of a series of blocks. We divided the series of blocks to ‘sequences’ of consecutive blocks. The defining characteristic of a sequence was that all its blocks shared the same target size. In Figs 4A,B, each point represent a block; the blocks connected by a line with a specific color represent a sequence.

A block consisted of 16 fixation trials. A fixation target was present at a fixed location throughout the trial. The target appeared in one of 8 possible locations, and the monkey had to maintain gaze within an invisible window around the target for a standard time interval. A successful trial was called a hit and the monkey was rewarded (see Methods for details). Crucially, if the target was too weak to be noticed one of two scenarios happened. Rarely, the eye happened to roam within the invisible window by chance and the trial was declared a hit. Usually, however, the trial ended up as an error. So the crucial result of a block was the count of how many of the block’s 16 trials ended up as hits.

As mentioned above, trials were run in sequences of blocks. Each sequence had a fixed target size; target size did vary from sequence to sequence. Each sequence is illustrated in Figs 4A,B with a specific color. In the first block of each sequence (the leftmost point in the appropriately colored trace) the target was so dim that the it went largely unnoticed. Indeed, the first blocks contain only 0-2 hits of the 16 trials, with the hits probably reflecting trials in which the eyes happened to be directed to the vicinity of the extremely weak target. The second block’s target was slightly more luminous, and then, for each subsequent block, the target’s luminance was slightly stepped up. After the first few blocks of a sequence, target luminance was high enough so that more of the block’s trials turned out to be hits; eventually, all, or almost all trials in the block were hits. At this level of performance, the target was evidently salient. More formally, we call the monkey’s performance in a block ‘full’, or ‘salient’, if there were 14-16 hits in the block’s 16 trials; ‘threshold’ if there were 3-13 hits. After completing a sequence, the monkey rested in the dark, allowing for refreshed dark adaptation, and then began another sequence, with smaller targets (usually with target radius being one half that of the most recently completed sequence). As targets became smaller, higher levels of luminance were needed to trace out the psychometric curve. Importantly, throughout the series of sequences, the entire range of scotopic luminance was monitored. The first sequences used very large targets (5.5 and 4 deg radius, correspondingly), and very dim lights were enough for the targets to be salient (just a little more than 10^−6^ cd/m^2^, in the low range of scotopic luminance range). For the smallest targets used, with 1/8 deg radii, only with luminance at the high end of the scotopic range (near 10^−3^ cd/m^2^) were the targets salient. Note how regular the psychometric curves look, even though they are based on quite few trials. Thus, these sessions track the threshold for saccades and fixation throughout the scotopic range.

To explore the monkeys’ performance, let us first ask, for each target size: with targets of that size, what level of luminance is needed for reaching threshold performance? What luminance level is needed for reaching salient, full performance? These questions are taken on in Figs 4C,E. Based on all the relevant sessions of each monkey, as expected, Fig 4C,E, show that the smaller the target the higher is the luminance needed, for both threshold and salience. The quotient between threshold and salience luminance levels remains close to 2 for all but the largest targets.

The regular relationship suggested by the graphs in Figs 4C,E of radius and luminance, for both threshold and salience performance levels, brings up the question whether a more general relationship holds. The number of photons over the entire target per unit time, technically proportional to the luminance intensity of the target, is a natural parameter to monitor. Figs 4D,F shows the data as a function of the target’s luminous intensity. For both threshold and salience performance levels, the total luminous intensity is almost fixed for small targets, with radius ≤ 1 deg (monkey M2), ≤ 2 deg (M3). The luminous intensity required for larger targets is higher. These findings are reminiscent of the psychophysical laws of stimulus detection, particularly of Ricco’s and Piper’s laws ^21^. They suggest that for small targets spatial summation is efficient, but for large targets summation starts to break up.

#### Upshift is higher near threshold than with salient targets

Let us now return to the upshift and ask whether the scotopic band setting is influenced by the task context. Namely, is the upshift contingent on the current block’s task context being salient or threshold? In addition, if we see an effect, the question will surface: could the effect reflect the same contingency as that on background luminance? Or could the contingency be of a new type?

Figs 5A,E show the mean upshift recorded in all relevant sessions of each monkey, in the threshold task described above. The colors of the points in Figs 5A,E encode the target sizes; these are the same colors as the psychometric curves Figs 4A,B. There are 2 solid lines in each of Figs 5A,E, thus 4 lines altogether. The circle-shaped points connected by the thin black lines encode targets at salience. These conditions are illustrated in the example block of Figs 4A,B as the dots above the top horizontal blue line. In these points the psychometric curves reach full performance. The square-shaped points connected by the thick gray lines encode targets at threshold performance. These threshold-performance data are illustrated in Figs 4A,B as the points between (and including) the two horizontal blue lines, between near-null performance to near-full. Thus, for each target size used, we get 2 points in Figs 5A,E: a circle-shaped point for the mean upshift at salient performance, and a square-shaped point for the mean upshift at threshold performance.

**Figure 5.**
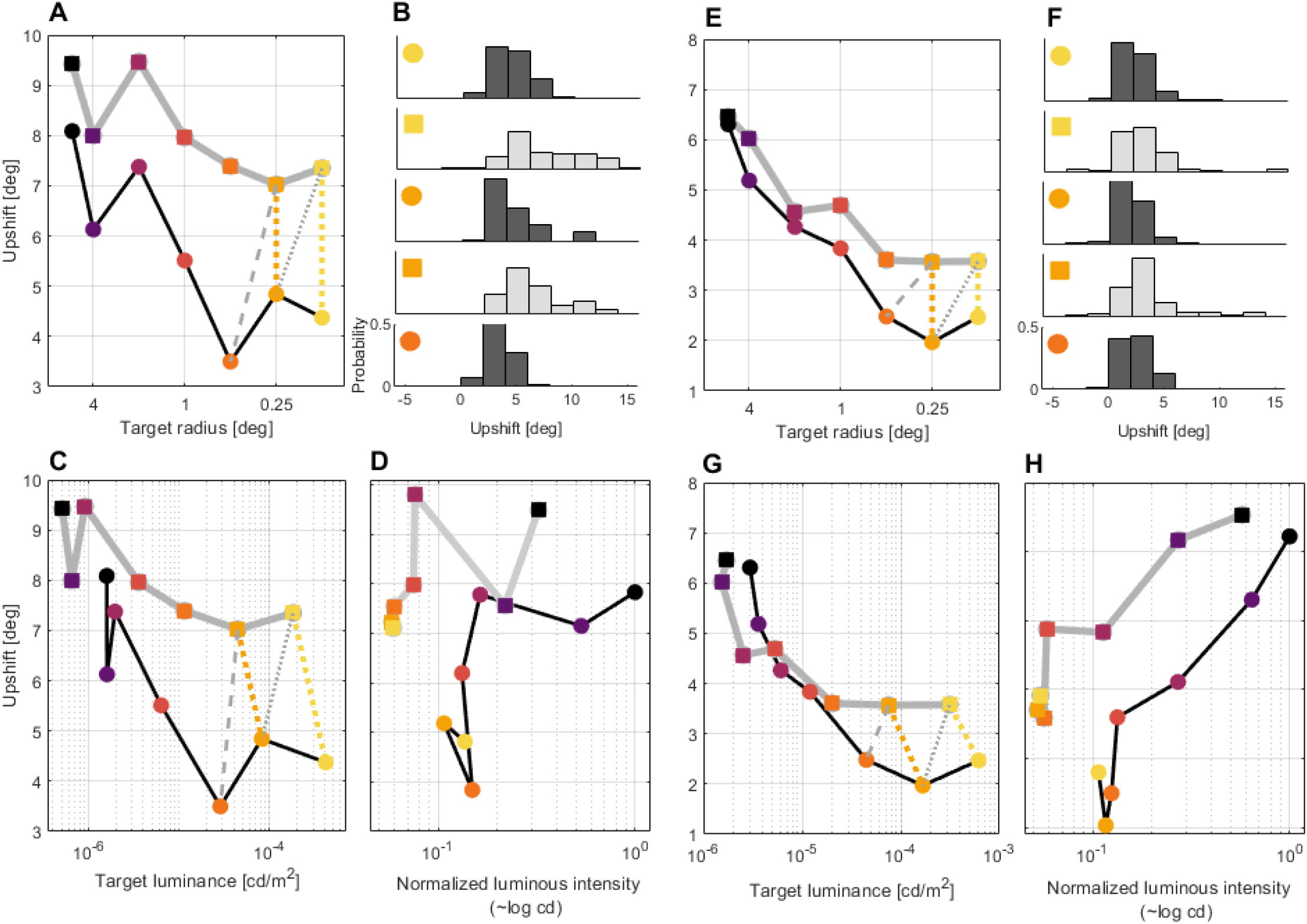
The scotopic center is set to more dorsal locations on the scotopic band when the task context is that of near threshold; this effect is not an automatic response to the lower luminance, nor luminous intensity. **(A)** The upshift (position of the scotopic center on the scotopic band) at threshold (gray line) and salience (black line) plotted as a function of target radius. **(B)** Histograms of the upshift at threshold (with the 2 smallest targets, gray bins) and salience (3 smallest targets, black bins). See markers together with (A) to identify the histogram. **(C)** Same data as (A), but plotted as a function of target luminance rather than target radius. **(D)** Same sata as (A), but plotted as a function of normalized luminous intensity. **(E) to (H)** Same format as (A) to (D), but data from monkey M2.

The main result of the threshold study now comes out. For targets of all sizes, for both monkeys, the mean upshift is higher at threshold than at salience. The scotopic band is set not in the same way in salient performance and near threshold.

Indeed, for all target sizes of both monkeys, the circle-shaped point lies below the square-shaped point. That is, the threshold performance is associated with higher upshift than salient performance; in other words, threshold is associated with more dorsal setting of the scotopic band. This is illustrated in Figs 5A,E explicitly for the two lowest-radius targets by the dotted yellow and bright orange lines that connect the circle and the square corresponding to each of the smallest targets.

The finding of the higher upshift at threshold than in salience is statistically very significant for the small targets, and less significant or insignificant for large targets. This qualification might reflect the less clear fixation task faced by the monkeys with large targets, and an interaction with upshift size. In more detail, for monkey M3, the upshift at threshold is significantly higher than at salience for all radii (paired t-test, *p<10*^*-4*^ for radius < 4 deg, *p<10*^*-2*^ for radius 4 deg and 5 deg). For monkey M2, whose mean upshift is smaller, this relationship is significant for small targets (*p<10*^*-3*^ for radii < 1 deg, and slightly significant or insignificant for larger radii (*p=0*.*02, 0*.*3, 0*.*06, 0*.*8* for radii 1, 2, 4, and 5.5 deg, respectively). Thus, for small targets the upshift is highly significantly larger at upshift than at salience. For larger targets, the mean upshift at threshold is higher, but less significantly or even insignificantly. Recall that the required gaze direction is less well defined for large targets, because gaze can direct to any part of the large target. This may lead to the compromised statistical significance in M2, whose upshift was lower. Indeed, in not one of these 4 target sizes was the mean upshift higher at salience than at threshold. In sum, the results support the observation that upshift is higher at threshold than at salience; a more dorsal location of the scotopic band appears to be used at threshold than at salience.

### Threshold or lower luminance?

The last section presented evidence that the upshift is greater at threshold performance than at salience. We set out to test upshift at threshold and at salience with the hope of showing that scotopic band setting is not always an automatic reflex-modification as in the response to background luminance (Fig 3). However, the observations we made could still, in principle, be automatic responses to increasing luminance; the only difference being that here the increasing luminance is of the target, not the background. By definition, at threshold targets of a fixed size are less luminous than at salience (otherwise there would be no problem seeing them). Does this observation explain away the observation of greater upshift at threshold performance level than at salience?

Let us consider a concrete example. Examining again Fig 5A, consider the rightmost, yellow circle and yellow square. The yellow circle shows that the mean upshift recorded while monkey M3 fixated salient targets of 1/8-deg radius was 4.37 deg. The yellow square shows M3’s mean upshift of the same targets, but at threshold performance, was 7.35 deg. The histograms of the upshift of individual trials in which M3 fixated salient 1/8-deg-radius targets is illustrated in the top panel of Fig 5B; the histogram of the threshold values is in the panel immediately below (see yellow square and circle markers on each panel). The histogram of the upshift of trials during salient performance occupy lower values than the histogram of the upshift of trials during threshold performance. It is thus unsurprising that the mean upshift at salience is significantly lower than the mean upshift at threshold (*p*<10^−6^). The yellow square and yellow circle in Fig 5A are connected to each other with a dashed yellow line. The line is vertical, as both the square and the circle reflect responses to targets of the same size, 1/8 deg. Now examine the same data in Fig 5C; here, the upshift is illustrated as a function of target luminance. The yellow vertical line is now tilted leftward, illustrating the fact that lower luminance is needed for threshold performance than for salient performance (compare Fig 5C to Fig 4C). This recapitulates the question: can we reject the null postulate, that the higher upshift of threshold performance only reflects the lower target luminance involved?

#### Threshold is not only lower luminance

Let us first add 2 points to the discussion. These are the points in Fig 5A,C that depict the upshift of M3 recorded while performing the ¼-deg sequence. Recall that the 1/8-deg points are the yellow circle and square, depicting the mean upshift in the salient and threshold blocks, respectively. The corresponding points depicting the mean upshift in the salient and threshold blocks are the bright orange circle and square, next to the yellow in Figs 5A,C. The bright orange dotted line connects the bright orange circle and square. In Fig 5A the bright orange line is vertical, as the yellow line, denoting that both bright-orange circle and square depict targets of the same size. In Fig 5C the bright-orange dotted line is tilted leftwards, just as the yellow dotted line, signifying that the ¼-deg threshold data was derived with lower luminance that the salience data. Fig 5B shows panels with the full trial-by-trial upshift histograms for the salience and threshold data of M3. Figs 5E-G show the analogous data for monkey M2.

So, is the upshift of 1/8-deg threshold (yellow squares in Figs 5A,C) higher than the upshift of 1/8-deg salience (yellow circles in Figs 5A,C) only because the 1/8-deg targets at salience are more luminous than the 1/8-deg targets at threshold? Let us now compare again the upshift of the 1/8-deg threshold (yellow squares) with salient performance, but this time with ¼-deg target (bright orange circle). The dotted grey line depicts this comparison in Figs 5A,C. In Fig 5A the dotted grey line is evidently tilted rightwards: this only reflects the fact that 1/8-deg radius targets are smaller than ¼-deg.

That the grey line tilts rightwards also in Fig 5C is much less self-evident. What this tilt reflects is the finding that the ¼-deg targets at salience are less luminous than the 1/8-deg targets at threshold.

Now consider what this finding means. At issue is whether the different upshift between salience and threshold only reflects the luminance of the relevant targets. We set out to check if the upshift at 1/8-deg threshold was higher than at 1/8-deg salience only because the threshold targets were dimmer. However, following this rationale, we expect the luminance relationship to generally reflect in the upshift size. Because the ¼-deg salience targets were dimmer than the 1/8-threshold targets, we expect the ¼-deg salience to entail a higher upshift than the 1/8-deg threshold. However, the result is inconsistent with this expectation. Fig 5B shows that the 1/8-deg threshold upshift histogram is at higher values than the ¼-deg threshold. The same situation is shown by M2 (Fig 5E-G). Hence the rationale at issue is rejected. The lower target luminance does not by itself explain the higher upshift associated with threshold, not for the case of the 1/8-deg threshold we followed in detail, hence not in general.

Thus, scotopic band setting appears to indeed be directly affected by the context of performing near threshold.

#### Not the target’s luminous intensity

As discussed earlier, Figs 4D,F showed that the target luminous intensity is very similar for targets of radius 1/8-deg to 1 deg (M2), and to 2 deg (M3). Only with larger radii is a higher level of luminous intensity needed. Thus, it appears that at least with small targets, reaching threshold requires about the same number of photos captured per second, invariant of the target’s size. This holds for both threshold and salient performance (each involves a different luminous intensity). Thus, the question comes up: could it be that the upshift reflects the luminous intensity of the target?

Figs 5D,H show the upshift as a function of the luminous intensity, for threshold (grey lines) and for salient performance (black lines). Focusing for a start on Monkey M3’s salient performance (circles connected by a thin black line in Fig 5D), the graph points corresponding to the performance with the small targets are arranged in a vertical line, reflecting the value of luminous intensity common to these small targets. The points corresponding to the small targets are spread out along this line segment. The same situation applies to the small targets in the threshold graph (squares connected by thick gray line in Fig 5D), as well as to the analogous graphs of monkey M2 (Fig 5H). Thus, the upshift changes while the luminous intensity remains almost unchanged. It follows that scotopic band setting does not reflect the luminous intensity.

The graphs of luminous intensity of the 2 monkeys at threshold (gray lines in Figs 5D,H) are positioned separately from the salience graphs. This separation between the graphs of threshold and salience suggests that a true mode change separates threshold performance from salience. The mode change involves moving to a different region of the luminous intensity – upshift space. Thus, in terms of luminous intensity, upshift at threshold might reflect a different mode of scotopic band setting from salience.

## DISCUSSION

This manuscript pursues the hypothesis that primate scotopic vision incorporates serial processing by a confined retinal region, ‘scotopic center’, much like the high-acuity processing by the fovea in photopic vision. Specialized sensorimotor transformations shift the scotopic center to the pertinent next location in the scene, and keep it there for processing the input. However, in difference from photopic vision, the scotopic center is not fixed on the retina, as is the fovea. Rather, there is a ‘scotopic band’. The scotopic center relocates on the scotopic band according to the ambient luminance and to the perceptual situation.

To test this hypothesis, we showed, first, that in scotopic darkness, a scotopic target is fixated by a confined retinal region dorsal to the fovea. We called this region the scotopic center. Second, centrifugal horizontal saccades shift the target’s image not to the fovea but directly to the scotopic center. These two points indicate that specific scotopic sensorimotor transformations enable the use of the scotopic center for scotopic serial processing. Third, persistent background scotopic luminance systematically relocates the scotopic center to a different position along the scotopic band. The darker the background is, the more dorsal is the scotopic center located on the scotopic band. Fourth, near-threshold conditions implicate a relocation of the scotopic center to a more dorsal position on the scotopic band. This scotopic band setting does not reflect only the physical characteristics of the target, specifically luminance and luminous intensity, and is likely to reflect the context of the task’s near-threshold conditions.

It is possible that scotopic band setting reflects other subtle processes. We alluded earlier to the bimodal histogram of the upshift of monkey M1 (see Fig 2A). The blue cluster suggested that M1 usually sets its scotopic band to high values. However, only in this monkey, a small subset of the trials showed no upshift (small blue cluster near the red cluster, near (0,0)). It is as if every now and then M1 cancelled the motor program for generating the upshift. This suggests the possibility that the upshift can be modulated by an even greater variety of non-automatic processes.

### Hypothesis on the function of the scotopic band

Our study is purely functional, but our eye-movement recordings intriguingly correlate to the previous histological results on the geometry of rod-density ^2,3,20,22^. Along the longitudinal axis of the band, going dorsally, rod density systematically increases whereas cone density decreases. We suggest that both the absolute value of rod density and the ratio of rod and cone densities are key parameters that determine visual sensation in the mesopic-scotopic range. Based on prior data, a position along the band is selected to best fit the expected input (that is, the scotopic band is set according to the expectations). We hypothesize that the same framework holds for mesopic vision, though the experimental data we present relates to scotopic vision. The predominant parameter for setting the scotopic band is the ambient level of light. In nature, ambient light changes slowly. The passage from full daylight through dusk to night darkness may take hours. Because the photon flux severely decreases, the information present in images changes, and the appropriate spot along the scotopic band changes respectively. However, ambient light is not the only factor necessitating scotopic band setting. An increase in sensitivity, even one caused by task constraints, also changes the scotopic band setting, as we have seen in the study of near-threshold scotopic vision. Thus scotopic band setting can happen automatically, reflecting the ambient light level, or in tandem with the pertinent cognitive context.

### Circuitry related to the scotopic band in the retina and in the brain

The local receptor density is the primary hard constraint limiting the perceivable information. From the receptors, information flows to the layers of the retina and on to the relevant regions of the brain. In the last few decades there was significant progress in understanding retinal information processing ^23^. Of particular relevance for our study is the discovery, in the mouse retina, that signals generated in rods may sometimes activate cone-based circuitry ^24^. This advanced level of understanding of information processing by retinas contrasts with the unawareness of the very existence in primates of a scotopic band. For all we know, the scotopic band might be limited to primates, in tandem with the fovea. The current manuscript’s description of the scotopic band opens up the question of the underlying retinal circuitry. If there is any similarity to the case of the fovea, connections of the receptors and the underlying retinal layers is bound to substantially differ in the scotopic band from the rest of the retina. Furthermore, this circuitry might gradually change along the longitudinal axis of the scotopic band, corresponding to the change in sensory processing. However, to gain any understanding as to what this circuitry is, anatomical and physiological studies must specifically target the scotopic band. Evidently, the relevant preparation is the monkey, both retina and brain.

What happens to information obtained by the scotopic band circuitry when it reaches the brain? This question is incredibly interesting too, and all open. Of special interest are the issues of the pertinent sensorimotor transformations, and their switching.

### Regional retinal specializations and nocturnal primates

Baden et al. recently reviewed the evolution of various aspect of vision across the 500 million years of evolution, in particular various configurations of receptor density, such as the fovea ^1^. The description of the scotopic band is pertinent to this important line of work. Very pertinent is also the description of preferred scotopic regions in the retinas of nocturnal monkeys. Wikler and Rakic ^2,20^ mapped the rod-density geometry in rhesus monkeys, owl monkeys, and bushbabies. In macaques, the maximal densities of cones and rods were, respectively, 160,000 and 180,000 receptors/mm^2^. In nocturnal owl monkeys and bushbabies cones were rare (less than 10,000 cones/mm^2^) but rods abundant (325,000 and 450,000 rods/mm^2^, respectively), with confined regions of high rod density present in both. This very high density reported in nocturnal primates does not only suport the contention that the scotopic band is indeed functional in scotopic vision; rather, these numbers beg the question of understanding whether there is serial scotopic processing also in owl monkey and bushbaby, and how the rod density specifically contributes to it.

### Situation in humans

Is the notion of the scotopic band relevant also for humans? The first paper that observed upshift in monkeys ^4^ reported that humans do not show upshift. Intriguingly, in terms of retinal anatomy, humans do show increased rod density dorsal to the fovea ^22^. A pertinent technical issue is that eye monitoring in the dark challenges the capacity of video-based eye monitors, because of the large changes in pupil size and shape. Perhaps, in human evolution, the motor program producing the upshift was partially or fully inactivated, or altogether erased. An interesting question to ask is whether this sensorimotor program enabling serial processing in scotopic vision was forever lost or can be reactivated.

## METHODS

### Subjects and surgical procedure

The monkeys of this study were used in other behavioral and neurophysiological experiments, and the general methods are described in other publications of the Thier laboratory in detail; see, in particular, (Dash et al., 2012; Caggiano et al., 2013, Khazali et al., 2017, Spivak et al., 2014). We used 3 *Macaca mulatta* monkeys; they are marked M1, M2, M3. All were male, 9-11 kg each. All of the monkeys were already trained in visual or oculomotor tasks. All experimental procedures are standard. In short, we used the scleral search coil method to record foveal gaze direction, also referred to as eye position. In the same procedure they were prepared for the eye position monitoring, the monkeys were prepared for neurophysiological recordings (not part of the present study). In particular, titanium head posts were implanted, allowing to robustly yet painlessly immobilize the heads during the experiment. Proper immobilization of the head is crucial for obtaining robust, accurate recordings of gaze direction. Surgeries were performed under intubation anesthesia with isoflurane and nitrous oxide, supplemented by continuous infusion of remifentanil (1 − 2.5 *μg*/(*kg* · *h*)). The relevant vital parameters were fully and tightly controlled throughout the procedure. All the procedures conformed to the National Institutes of Health Guide for Care and Use of Laboratory Animals and were approved by the local ethical committee (Regierungspräsidium Tübingen).

### Setup

The experiments were performed in a completely lightproof electrophysiological setup. The monkeys were seated in front of a cathode ray tube (CRT) screen, at a distance of 35 cm from the screen center. The CRT was an Eizo Flexscan F730, 50-cm diagonal, displaying 1024×768 pixels at a frame rate of 60 Hz. The only source of light during the experiment was the CRT monitor. Gaze direction (eye position) was sampled at 1000 samples per second.

The experiments were run, and data were collected, using the open source measurement system nrec, https://nrec.neurologie.uni-tuebingen.de/nrec/nrec, created by F. Bunjes, J. Gukelberger and others, and used as standard in the Thier lab at Tübingen. The nrec output files were converted to Matlab (https://www.mathworks.com/, MathWorks, Natick, Massachusetts, USA), and post hoc data analysis was carried out in Matlab.

### Studies

A session comprised the data collected in a single day. There were 3 studies, each having its trial and session structure. *Study 1* (Fig2): fixation and saccade in scotopic dark with small scotopic-bright targets (revealing the scotopic center); *study2* (Fig 3): dependence of the upshift on background luminance (revealing the scotopic band); and *study 3* (Fig 4): dependence of the upshift on near-threshold. We now describe the stimuli, trial and session structure in these studies.

We first ran study 1 on monkeys M1,M2,M3. Then we ran study 2, on M3 and M2. Finally we ran study 3, again on M3 and M2. Because study 3 substantially differs in procedures from studies 1 and 2, we first describe the methods of studies 1,2, and then of study 3, separately.

### Studies 1 and 2

#### Visual stimuli

The CRT’s background was always uniform and featureless. In photopic conditions, the background luminance was 7 cd/m^2^. In scotopic conditions it was dark, less than the scotopic threshold.

In the studies 1 and 2, the targets were circular, uniformly filled. In the study 1, photopic targets were small and bright (0.02 deg radius, 60 cd/m^2^). Scotopic targets were slightly larger and dimmer (0.14 deg diameter, 7*10^−4^ cd/m^2^). Thus the scotopic targets were bright for scotopic conditions, but well within the scotopic range. The central fixation spot and peripheral saccade targets were identical (except for position and time in trial). In the first part of study 2 (Fig 3A) the targets were small (0.02 deg radius) and bright (60 cd/m^2^), similar to values we previously used to study photopic upshift ^6,9,25^; whereas the background luminance in the first part of study 2 (Fig 3A) changed from block to block, stepping from scotopic to photopic: 10^−6^ cd/m2 (blue markers), 10^−5^ cd/m2 (yellow markers), 10^−4^ cd/m2 (cyan markers), 10^−3^ cd/m2 (black markers), 10^−2^ cd/m2 (green markers), 7 cd/m2 (red markers). In the second part of study 2 target luminance was 10^−3^ cd/m2 and target radius: 0.25 deg. Background luminance differed from block to block, from scotopic to photopic: dark (blue marker), 0.4*10^−5^ (green marker), 10^−5^ (cyan marker), 0.2*10^−4^ (grey marker), 0.4*10^−4^ (orange marker), 10^−4^ (purple marker), 2*10^−4^ cd/m^2^ (yellow marker).

#### Locations used for targets

Study 1: in a daily session we used a set of 18 or 20 target locations, spaced 2 deg away from each other, all positioned on the horizontal meridian. In order to increase the total range of saccade sizes (for aims beyond the present study), target locations were slightly shifted from day to day. The closest targets to the central fixation spot were positioned at ±1 deg, ±1.5 deg, ±2 deg, or ±2.5 deg from the central fixation spot. Thus, the maximal horizontal eccentricity was 20.5 deg. Here we pulled together all data, taking into account the target location of each individual trial, regardless of the session’s target set.

Study 2: the fixation targets had three possible locations, positioned on the horizontal meridian: the center of the screen, 10 deg to the right and 10 deg to the left of the center. Every block consisted of 30 trials; every target location was used 10 times in a block.

#### Trial structure

Study 1: when the central fixation spot appeared, the monkey was required to direct his gaze into an invisible window around the spot within 1 s (see next paragraph for detail). Then the monkey had to maintain fixation, that is, to keep his gaze within the window, for at least 1.5 s. Next, the central fixation spot disappeared, and, simultaneously, the peripheral target appeared. The monkey had to make a saccade into an invisible window around the peripheral target within 1 s. The target remained for 2 s on the screen and the monkey was required to keep his gaze within the invisible window around the peripheral target until the end of the trial. On detection of an error, the computer aborted the trial and initiated a new trial; on correct performance (‘hit’) the computer rewarded the monkey with a drop of water. Total trial duration was 4.5 s.

Study 2: the monkeys had to direct their gaze into the invisible window around the target spot, and keep their gaze there as long as the target remained on. The fixation invisible window appeared 1 s after the appearance of the target. Thus, within 1 s from target onset the monkey had to shift his gaze into the invisible window around the target. The trial was counted as a hit if the monkey maintained his gaze within the invisible window centered on the peripheral target for an additional 1.5 seconds. At the end of the trial, the monkey was rewarded for performing a hit with a drop of water as reward. After the completion of the trial, the next trial immediately followed. The location of the target changed from trial to trial in pseudo-randomized order. The total duration of the trial was 2.5 seconds.

#### Gaze direction windows

In photopic conditions, the invisible windows were square shaped, with 3 deg radius (±3 deg for each horizontal and vertical). In scotopic conditions (including the mesopic blocks in study 2), the windows were 15 deg vertical radius and 5 deg horizontal radius. These large invisible windows reflected our wish not to artificially constraint the upshift. For details see ^2–4^.

Does the large scotopic invisible window allow noise into the fixation data?This choice of parameters reflects our experience with studying the upshift. As can be seen in all the panels of Figs 2A, and in the previous studies of the upshift for photopic upshift, the gaze positions densely accumulate in a small sub-region of the large window. Fig 3 shows an example of how the clusters move upon appropriately changing the visual conditions. These clusters (and others observed in previous studies) show that the large windows allow the physiological changes in gaze direction to come through, rather than increase noise.

#### Daily sessions

A daily experimental session consisted of a series of blocks of trials, 100 (study 1), 30 (study 2) trials in each block, counting only hits. A session began with standard gaze direction calibration, in photopic conditions; the calibration was followed by a block or two of photopic trials. Then the monkey waited through a 45-min interval of dark adaptation. During this time, the monkey was in full darkness, and did not work. Then the scotopic blocks were recorded.

#### Data analysis

Study 1: to calculate the mean gaze direction during central and peripheral fixation (Fig 2 A, B) we averaged across intervals of 250 ms, one interval before the saccade started, and one after the saccade was completed. For the fixation of the central spot, the interval was sampled before the central fixation spot was turned off. For the fixation of the peripheral target, the interval was sampled at the end of the post-saccadic target fixation, shortly before the end of the trial. Therefore, the peripheral target fixation measurement was not contaminated with saccadic correction.

Study 2: Mean gaze direction during target fixation was calculated by averaging across the last 1 s fixation of the trial.

The parameters of the study 3 described in Fig 4 are explained in a separate section, ahead.

### Study 3

Study 3 aimed at studying the upshift near threshold, using targets of different sizes. The experiment was conducted in scotopic conditions that required 45 minutes of dark adaptation prior the task. The standard gaze direction calibration was made in photopic conditions before the dark adaptation. Every trial began with an appearance of the fixation target at one of the eight locations arranged in a circle of 15 deg radius. Targets were presented in a pseudo-random order, one target per trial. A target remained on the screen for two seconds without fixation window around it. This time was intended to allow the monkey to find target location and stabilize gaze at it. If at the end of the two seconds the monkey still gazed at the target, fixation window appeared for another 0.5 seconds. Fixation of the target during the additional 0.5 seconds was rewarded with a drop of water, and the trial was labeled as correct. If the monkey broke fixation during the last 0.5 seconds of the trial or did not look at the target before the fixation window appeared, the trial was labeled as incorrect and immediately switched to the next. No reward was given in this case. Possible trial length was 2-2.5 seconds depending on monkey’s performance. Two seconds trial length indicates that the monkey did not locate the target. Trial length of 2.5 seconds indicates that the monkey detected the target and completed the trial successfully. This trial was counted as hit. Any trial length between 2 and 2.5 seconds would suggest that the monkey detected the target correctly but broke fixation during the last 0.5 seconds of the trial. In this case the trial was not counted as hit. Every block consisted of 16 trials, so that every target appeared 2 times in a block. In every session we ran several blocks. It allowed us to construct psychometric curves (Fig 4) taking into account the correct trial number and target luminosity in every block. The first block in a session contained targets at very dim luminosity that as we knew the monkey could not detect. We increased target luminosity from one block to the next by the constant range of units that was determined by the stimuli-displaying software. By increasing target luminosity from block to block we also increased the ability of the monkey to detect the target. The session was accomplished when the monkey could complete all 16 trials in a block successfully. The monkey underwent additional dark adaptation of 10-15 minutes between the sessions. The possible target radii were: 2.4, 2, 1, 0, -1, -2, -3, -4 and -5 in log^2^ degrees. The largest target’s radius was set to 2.4 instead of 3 log^2^ degrees due to monitor’s limitations. The size of the fixation window was 15 deg horizontal and 20 deg vertical. This size ensured that even the largest fixation target could fit the window. Target’s radius decreased from session to session in a consecutive order. At every session only one target radius was used. Mean gaze direction during target fixation of the correct trial was calculated by averaging across the last 0.5 s fixation of the trial.

